# Heat stress conditions affect the social network structure of free-ranging sheep

**DOI:** 10.1101/2023.02.26.530064

**Authors:** Zachary Borthwick, Katrin Quiring, Simon C. Griffith, Stephan T. Leu

## Abstract

Extreme weather conditions, like heatwave events, are becoming more frequent with climate change. Animals often modify their behaviour to cope with environmental changes and extremes. If the environmental conditions influence the trade-off between an individual’s social propensity and optimal thermoregulation through shade use for instance, then divergent social decisions may be made. Hence, such behavioural changes have the potential to influence an individual’s position in its social network, and the social network structure as a whole. We investigated whether heat stress conditions (quantified through the Temperature Humidity Index) and the resulting use of shaded areas, influences the social network structure and an individual’s position in it. We studied this in free-ranging sheep in the arid zone of Australia, GPS-tracking all 48 individuals in a flock. When heat stress conditions worsened, individuals spent more time in the shade and the network was more connected (higher density) and less modular. Furthermore, an individual’s social connectivity scaled with its shade use behaviour. Interestingly, individuals with intermediate shade use were most strongly connected (degree, strength, betweeness), indicating their importance for the connectivity of the social network during heat stress conditions. Our study shows that heat stress conditions, which are predicted to increase in severity and frequency due to climate change, influence the resource use within the ecological environment. This has flow on effects for the animal’s social environment through the changed social network structure.

**Significance statement:** Due to climate change, animals experience heat stress conditions more frequently. When ambient temperatures are high, individuals can seek shaded areas to mitigate the effect. However, whether these behavioural changes affect the interaction patterns in social species is not well understood. We tracked all individuals in a flock of sheep using GPS collars. We demonstrate that when heat stress conditions worsened (higher temperature humidity index), individuals spent more time in the shade and the social network was more connected with less internal structure. Furthermore, an individual’s social connectivity scaled with its shade use behaviour, and individuals with intermediate shade use were most strongly connected in their social network. Our findings illustrate that individuals respond to climate change induced environmental conditions and that this has flow on effects for their social connectivity and the social network of the group.

## Introduction

The climate is changing, resulting in a greater frequency and severity of heat stress conditions (Sejian et al. 2018). These conditions can have several effects on individual animals, including impaired endocrine and immune function (Mete et al. 2012; Lacetera 2019) as well as reduced reproductive success (van Wettere et al. 2021). Heat stress also elicits changes in behaviour to avoid hyperthermia. Individuals can limit or change their activity patterns when ambient conditions are particularly challenging (Bourgoin et al. 2011; Hetem et al. 2012; Fuller et al. 2016; Leu et al. 2021). Similarly, animals can avoid microclimates that are hot and in direct sun and shift their space use to favour shaded areas (Hetem et al. 2012; Leu et al. 2021). Reduced food intake to decrease metabolic heat production, increased drinking frequency, and increased standing time to increase surface area for heat loss by convection (Galán et al. 2018) have also been recorded during hot conditions. Furthermore, how strongly individuals respond to changing ecological conditions can depend on individual attributes such as genetic and phenotypic traits, metabolic status (Galán et al. 2018), sex (Li et al. 2018), and body condition (Parer 1963).

Behavioural responses to heat stress also reflect a cost-benefit trade-off. For example, bats (*Myotis yumanensis, Antrozous pallidus* and *Tadarida brasiliensis*) trade off thermoregulation opportunity with safety when selecting roosting sites (Licht and Leitner 1967), and alpine ibex (*Capra ibex*) trade off thermoregulation and foraging opportunities (Mason et al. 2017). These and other behavioural changes are considered the first response of individuals to heat stress (Sejian et al. 2018) and can increase the chances of survival (Li et al. 2018). However, thermal tolerance varies among individuals (Drown et al. 2021) and hence individuals have to trade off optimal thermoregulation with their social propensity. Here, we investigated how individual thermoregulatory behaviour in a group of free-ranging sheep in the arid zone of Australia affects individual social connectivity as well as group social network patterns.

During heat stress conditions sheep spend time in the shade as a behavioural response to support physiological thermoregulation (Leu et al. 2021). Sheep also spend more time at watering points to avoid dehydration, and activity levels are substantially lower compared to more typical conditions (Leu et al. 2021). Sheep that were kept under tree shade showed fewer heat stress related behaviours and shade use behaviour reduced radiant heat load (De et al. 2020). Furthermore, individual sheep are repeatable in their shade use behaviour, but the length of time spent in the shade varies among individuals (Leu et al. 2021). We hypothesise that behavioural responses to heat stress conditions, in particular shade use, affect individual social association patterns.

Furthermore, the social structure of a group reflects individual behaviour that maximises fitness (Emlen and Oring 1977), and we further hypothesise that these behavioural responses upscale to the group level and affect the social structure of the study group. Two alternative effects on the network are conceivable. Sheep could aggregate in a few large, shaded areas. This would result in social networks that are more connected (higher density) and are less structured (lower network modularity). If individuals interact with conspecifics that are present in shade without social preference, this will lead to homogenised association patterns (low coefficient of variation of edge weights). Alternatively, if animals use several different shaded areas, networks are disrupted and become less connected. This would lead to lower network density, greater modularity and heterogeneous interaction patterns, i.e. higher coefficient of variation of edge weights. We refrain from making a priori predictions as available shade was not limited but we do not know whether sheep would aggregate in a few or many separate shade patches.

Group coordination and synchronised action is important in social species affecting social network structure (Maldonado-Chaparro et al. 2018). Social network structure in turn affects population processes such as social information (Aplin et al. 2012) and parasite transmission (Sah et al. 2017; Leu et al. 2020), as well as mating/reproductive success (Beck et al. 2021). Understanding how behavioural responses to heat stress conditions affect an individual’s position in its social network and the social network structure of the group will provide important insight how climate change may affect population processes of social animal species.

## Materials and Methods

### Study site and tracking of sheep

We conducted this study at Fowler’s Gap Arid Zone Research Station (31◦05′ S, 141◦43′ E) in summer 2018. We observed 48 female sheep in a fenced paddock of 6 × 1 km for 82 days. All individuals were non-pregnant female Merinos, born in mid-2016 and approximately 1.5 years old. Each individual was equipped with a GPS collar (Global Position System, i-gotU GT-120 by MobileAction, with a larger battery CE04381 by Core Electronics). The GPS collars recorded positions every 2 minutes. The GPS collars were synchronised and recorded the location of all sheep at the same time. The collars weighed 700 g or 1.8 % of the mean body mass of our sheep (39.3 ± SE 0.7 kg) and did not impact sheep movement behaviour (Leu et al. 2021).

### GPS Data Processing

We followed the same data processing procedure as in Leu et al (2021). In brief, spatial outliers were removed from the raw GPS data using three methods, (1) all locations that used less than 3 satellites or were clearly outside the fenced study area; (2) locations that an individual could not have moved to at a maximal movement speed (180 m per 2 min i.e. 5.34 km/h; Manning et al. 2014) and (3) locations that resulted in spikes away from the movement path, filtered based on maximal speed and large turning angles. Because GPS data loggers recorded locations at slightly different times, we interpolated each individual’s filtered data to exactly the same two-minute intervals.

### Inferring sheep social networks from spatial locations

We conducted all social network construction and statistical analysis using R statistical software (R Core Team 2021). First, we considered two individuals to be associated if they were within one body length (approx. 1 m) of one another at the same time. Then, taking the precision (2m) of both GPS units into account, this leads to an association threshold of 5m (body length 1m + precision 2 × 2m = 5m). Using this threshold, we identified all associations among all possible pairs of individuals and constructed a social network for the midday period of each study day (n=82). We focussed on the midday period with 1.5 hours before and after the zenith of the sun (3 hour period), as radiation is highest and shade use would be the most beneficial at this time (Leu et al. 2021).

We used the R package spatsoc (Robitaille et al. 2019) to apply the spatio-temporal criteria and generated a group by individual matrix (GBI) for all dyads. Then, using the package asnipe (Farine 2013), we generated an individual by individual adjacency matrix and calculated the simple ratio index as edge weight. The edge weight reflects the proportion of time a dyad spent in close contact relative to the time they were both recorded.

For each midday network, we calculated three whole network measures – network density, modularity and coefficient of variation of edge weights. We then related these measures to the temperature humidity index (see below) which is a proxy for the ambient heat conditions. We have previously shown that network density can be influenced by environmental characteristics such as habitat structural complexity (Leu et al. 2016). Here, we were interested whether shaded areas and their use influences the network. We used the package igraph (Csárdi and Nepusz 2006) to calculate the network density for each midday network. Network density is the ratio between existing links and all potential links of a network. A high value of network density indicates a highly connected network (Sosa et al. 2021). Second, we calculated modularity as it gives insight into the internal structure of the social network and captures the result of fission-fusion events (Sosa et al. 2021) which could take place due to different shade use when faced with heat stress conditions. A high measure of modularity indicates the presence of smaller subgroups that frequently interact with members of the same subgroup but interact less frequently with members of other subgroups. Third, we calculated the coefficient of variation of edge weights. The coefficient indicates how heterogeneous dyadic relationships (length of time spent together) were across the social network. A greater coefficient of variance indicates greater heterogeneity, that is some pairs spent more time together than other pairs. This can either be few but long periods, or many short periods. This metric is particularly informative alongside density and modularity. Density and modularity inform about the structure of the whole network, while the coefficient of variance of edge weights informs how structured individual associations are.

In addition to the whole network level, we also investigated the effect of heat stress conditions at the individual level. For each individual, in each 3-hour midday network, we calculated three node-level metrics, degree, strength, and betweenness centrality. We related these measures to individual shade use behaviour. Degree represents the number of individuals an animal associates with during the given time period, strength is the sum of edge weights of the node (Sosa et al. 2021). Betweenness centrality measures the number of times a node is included in the shortest paths between two other nodes (Sosa et al. 2021). Hence, it reflects the extent to which an individual connects sub-groups. It indicates the relative importance of an individual for transmission processes, e.g. transmission of information or pathogens (Newman 2005), which have been shown to have fitness consequences (Gilby et al. 2013).

### Calculating Temperature Humidity Index (THI) and individual shade use

Different Bioclimatic Indices (BCI) exist to assess the impact of climatic conditions on livestock reared outdoors (Theusme et al. 2021). In this study we used the Temperature Humidity Index (THI) which is a suitable climatic parameter to describe heat stress conditions in sheep (Marai et al. 2007; Theusme et al. 2021). It is calculated as THI _sheep_ = T – {0.31 – (0.31 x (RH/100))) x (T – 14.4)}; where T= Ambient Temperature (in °C) and RH = Relative Humidity (in %). The dry bulb temperature and relative humidity were available every hour from a nearby Bureau of Meteorology Automated Weather Station (Fowlers Gap field station AWS41628). Hence, we calculated the mean THI for the 3-hour midday period around the zenith of each day. In our analysis we used THI as a continuous variable. Nevertheless, different THI thresholds have been identified to indicate different levels of heat stress for sheep; no heat stress < 22.2 THI, moderate heat stress 22.2 to < 23.3, severe heat stress 23.3 to < 25.6 and extreme severe heat stress ≥ 25.6 (Marai et al. 2007).

We also calculated how long each individual sheep spent in the shade during the 3-hour midday period of each day. We considered that trees with a minimum height of 1 metre provide shade for a standing sheep. We mapped 106 tree patches using a handheld GPS unit (Leu et al. 2021). Each tree patch was a georeferenced polygon with known boundary coordinates.

### THI and shade use effects on network structure

We used three separate linear models to investigate the relationship between THI (continuous variable) and the whole network metrics (*density, modularity and coefficient of variation of edge weights* – all continuous variables). We then determined the relationship between THI (continuous) and individual shade use behaviour (minutes in shade per 3-hour midday period, continuous) using a linear mixed model. Time spent in shade was the dependent variable, THI the independent variable and sheep *ID* (48 levels) a random effect. Finally, we investigated whether shade use behaviour affected an individual’s social network connectedness. We scaled and centred the shade use variable using the ‘scale’ function in base R (R Core Team 2021) to meet model assumptions. We used three separate linear mixed models. *The models with strength* and *betweenness centrality* (both continuous) as the dependent variable used a Gaussian distribution, whereas the model for *degree* (discrete count data) used a Poisson distribution. We included *shade use* and (*shade use)*^*2*^ both as fixed effects (continuous) as this better followed the data structure and improved model fit. As before, sheep *ID* (48 levels) was a random effect.

### Permutation based hypothesis testing

Social network measures are inherently non-independent and therefore violate an important assumption of most statistical tests (Croft et al. 2011). Combining linear models with permutations accounts for the non-independence and also allows to identify the effect of the investigated variable beyond other (unmeasured) variables that contribute to the social structure (Farine 2017). We created 1000 random social networks for each analysis, deduced the same network measures and reran the same linear models. We then compared the model coefficient based on the empirical social network with the distribution of coefficients derived from the random networks. Two-sided-p-values for each effect were calculated as twice the number of times a randomised coefficient was more extreme than the observed test statistic divided by 1000, the number of randomisations (Farine 2017).

We conducted pre-network permutations when investigating whole network metrics. We used the ‘network_permutation’ function in asnipe (Farine 2013). This involved carrying out 1000 randomisations of the group-by-individual (GBI) matrices, which assigned individuals to different, random dyads. Using these randomised GBI matrices, we generated 1000 randomised adjacency matrices and deduced the relevant network measures. We conducted node label randomisations when investigating node-level metrics. We used the ‘perm.net.nl’ function in the ANTs package (Sebastian et al. 2018). We used shade use as node labels and directly randomised the shade use values to generate 1000 shuffled data sets for the midday period of each day.

## Results

### Heat stress conditions and shade use

Across the 82 days of the study period, individuals experienced on average a temperature humidity index of 28.9 (SD = 3.5) with a min of 20.72 and max of 34.29. Most days (81.71%) fell into the extreme severe heat stress category, followed by 10.98% in severe heat stress, 4.88% in moderate heat stress and 2.44% in the no heat stress category. The average tree height was 2.23 metre and 84 of the 106 shaded areas were used by our sheep. Time spent in the shade increased with an increase in heat stress conditions (THI). During the midday period (180 min) sheep spent a mean of 66.9 minutes in the shade when conditions were categorised as extreme severe heat stress compared to only 20.8 minutes when conditions were classified as no heat stress.

### Heat stress conditions and shade use affect network structure

Network density increased with THI, whereas modularity and coefficient of variation of edge weights decreased (Table 1).

**Table 1.** Effects of Temperature Humidity Index (THI) on whole network structure

| Dependent variable | Coefficient | SE | t | R<sup>2</sup> | P<sub>rand</sub> |
| --- | --- | --- | --- | --- | --- |
| Network Density | 0.02 | 0.005 | 3.82 | 0.14 | <0.001 |
| Modularity | -0.014 | 0.005 | -2.699 | 0.07 | <0.001 |
| CV of Edge Weights | -0.14 | 0.03 | -5.09 | 0.24 | <0.001 |

Individual shade use behaviour increased with THI (estimate = 6.88, SE = 0.26, t-value = 26.61, R^2^=0.16, p<0.001). Shade use was associated with all three node-level social network measures. More specifically, the effect of shade use and (shade use)^2^ was comparable for degree, strength and betweenness centrality (Table 2). The combination of the positive coefficient for shade use and negative coefficient for (shade use)^2^ means that as shade use increased, sheep showed greater social connectedness, which then decreased again when individuals spent even longer periods in the shade (Fig 1).

**Table 2.**
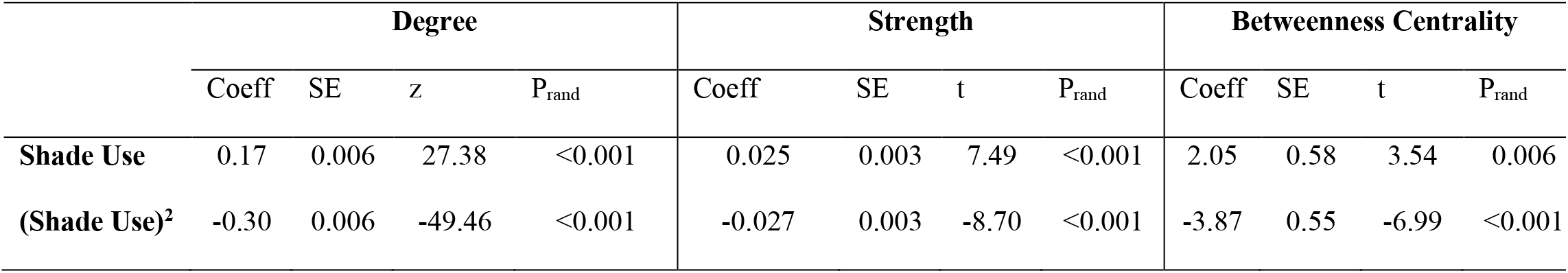
Effect of Shade Use and (Shade Use)^2^ on node-level network metrics

**Figure 1.**
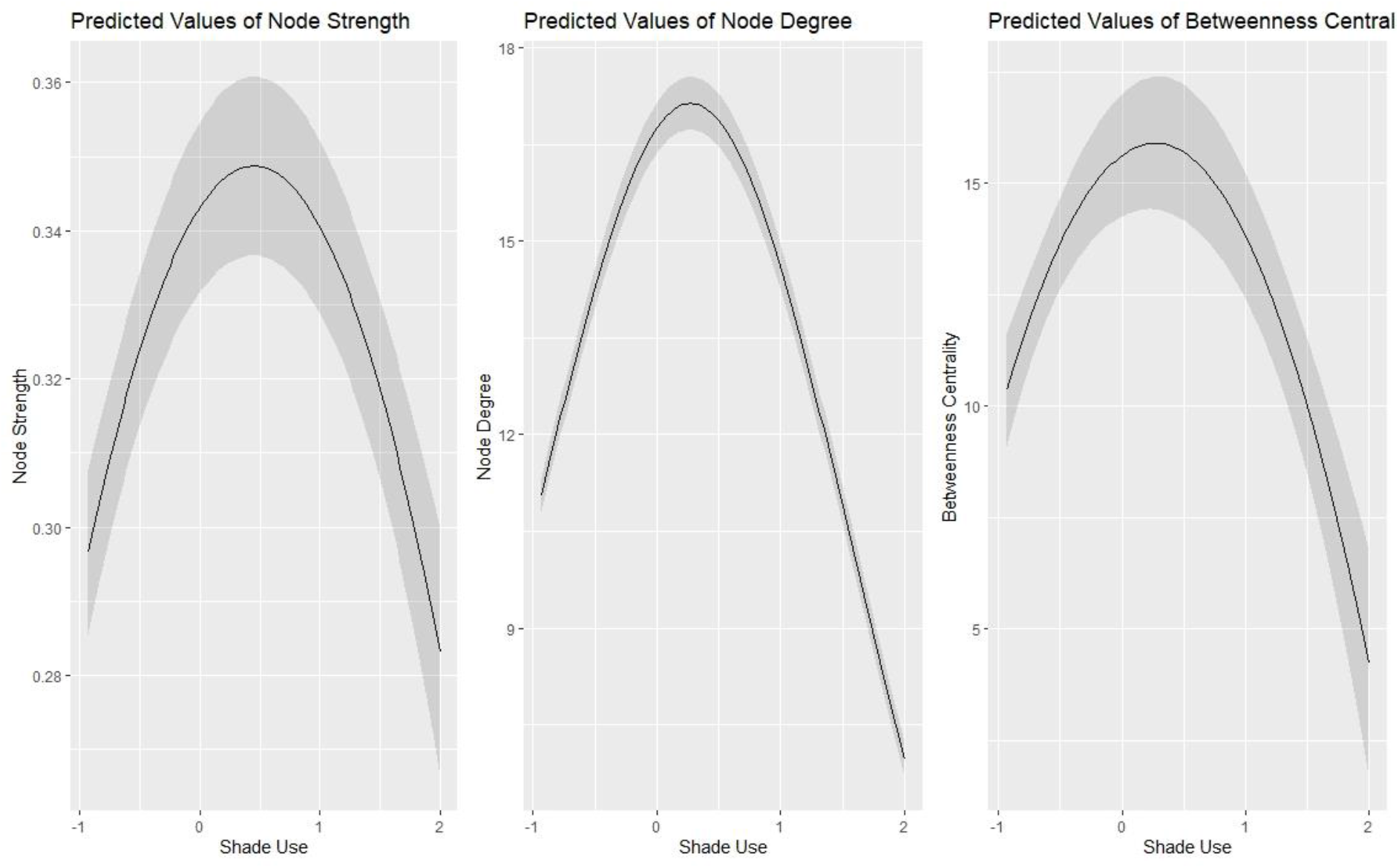
Estimated values of node strength, degree and betweenness centrality against shade use (scaled and centred) are depicted in three separate graphs.

## Discussion

### Heat stress conditions affect whole network structure

As heat stress conditions worsened, indicated by an increasing temperature humidity index (THI), networks became denser, less modular and the animals’ social associations became less heterogeneous (decrease in CV of edge weights). That is, the differentiation into stronger and weaker social ties was reduced and social associations were more similar. Taken together, this suggests that with increasing heat stress conditions, the social network was more connected but lost internal structure. Previous studies have shown that external stressors can impact social network structure. Maldonado-Chaparro et al. (2018) showed that even temporary disruptions, such as splitting a group of zebra finches into sub-groups for two days affected the social network after the sub-groups were joined again. However, in contrast to our study, the disturbance effect of group splits increased the internal group structure and caused more differentiated social relationships. This further reduced foraging efficiency as fewer individuals accessed the foraging patches at the same time (Maldonado-Chaparro et al. 2018). The more connected and less structured network in our study could lead to improved transmission processes (Sah et al. 2017) which can be beneficial or costly depending on the transmitted agent. Improved information transmission can increase the capacity of individuals to find resources such as food or water (Aplin et al. 2012; Kendal et al. 2015) and in our study, shade. Furthermore, predator detection could also be improved (Frechette et al. 2014). For example, although aggregations are more vulnerable to being detected by predators (Pulliam 1973), faster information spread about the presence of predators could provide protection from actual predation. Furthermore, in the context of this study, shaded areas are likely low-risk as they provide cover, which reduces detection by predators and reduces glare that could hamper the prey’s ability to monitor its environment (Carr and Lima 2014; Sabal et al. 2021).

However, a well-connected social network with low modularity also improves the spread of pathogens and parasites which has negative well-being and health consequences (Griffin and Nunn 2012; Nunn et al. 2015). Furthermore, disease transmission may be particularly detrimental when animals are already heat stressed, as stress has been shown to impair immune system function (Hing et al. 2016). Hence, the costs of a connected and homogeneous network structure could outweigh the benefits, even if heat stress conditions are rare and short lived. This is because pathogen transmission processes often only require a few short interactions. However, the cost benefit ratio of highly connected networks in sheep remains to be investigated.

### Shade use behaviour affects individual network connectivity

Sheep varied in the extent of their shade use behaviour. During periods of extreme severe heat, time spent in the shade ranged from 0 to 180 minutes over the 3-hour period. We previously showed that shade use behaviour varies among individuals, but that individuals are consistent in their own behaviour between heat wave events (Leu et al. 2021). Competition among individuals, which can be further modulated by the social hierarchy, can influence access to resources (Nelson-Flower et al. 2011; Majolo et al. 2012). However, this is not the case in all species. In the sociable weaver (*Philetairus socius*) for instance, which lives in complex cooperative societies, shade use behaviour, and hence access to shade, was not influenced by social rank (Cunningham et al. 2017).

Nevertheless, dominant social weaver individuals could maintain their body temperature more precisely at increasing ambient temperatures, whereas subordinate individual could not (Cunningham et al. 2017). At this stage we do not know whether social hierarchy plays a role in the shade use behaviour in sheep. For example, whether high ranking individuals restrict access to certain patches of shade. This was beyond the scope of the current study. Instead, we were interested in how the variation in using shade patches affects social connectivity in sheep.

Shade use had a non-linear relationship with an individual’s social connectivity. First, social connectivity increased with shade use behaviour, before reaching a peak and then decreasing again. This was the case for node degree, strength and betweenness. The initial increase in social connectedness may reflect fusion events with some sub-groups aggregating at certain shade patches. Our paddock included 106 tree patches, 84 of those patches were used during the study period, which indicates that shaded areas were unlikely limited. This contrasts with other studies that showed an increase in node strength in response to ambient conditions. For instance, the increase in node strength in networks of fairy wrens (*Malurus melanocephalus*) was due to individuals flocking to limited habitats after a disruptive fire (Lantz and Karubian 2017).

Interestingly, individual sheep with intermediate shade use behaviour were most strongly connected in the social network. Whereas individuals that spent either less or more time in the shade were less connected. One possible explanation is, that those intermediate individuals spent some time in one shade patch, then moved away into the open where they may encounter and interact with individuals that remained active and spent little time in the shade. This is then followed by another period in the shade, either in the same or a different shade patch. Furthermore, sheep with intermediate shade use behaviour appear to play an important role in the overall social network connectivity. This notion is supported by our finding that intermediate shade users also had the highest betweenness values. Highly connected individuals (high degree and strength) in important network positions (high betweenness) play a crucial role for maintaining network cohesion, stability and for social transmission processes (Wey et al. 2008) For instance, in pigtailed macaques (*Macaca nemestrina*), a small subset of well-connected individuals significantly contributes to maintaining stable social environments during periods of chronic perturbations. Without these individuals, group members build smaller, less diverse, and less integrated networks (Flack et al. 2006). In such situations where some individuals are responsible for maintaining links between groups that are otherwise less connected, weak ties emerge which can serve as transmission points (Vanderwaal et al. 2016). As mentioned above this could be beneficial for the population if information is transmitted or costly if parasites or pathogens are transmitted. Irrespective of the agent that is transmitted, it shows the disproportionate influence of these individuals for population processes.

In conclusion, heat stress conditions affected shade use behaviour in sheep, their individual social connectedness and group social network structure. Importantly, individuals with intermediate shade use behaviour occupied well-connected and important network positions. This suggests that they play an important role in maintaining social network cohesion and function during periods of heat stress. Our study shows that climate change which is predicted to lead to more frequent and more severe heat stress conditions, may not only impact the ecological environment that individuals experience but also their social environment through changes of the social network structure.

## Acknowledgements

We thank Keith Leggett (Director Arid Zone Research Station, Fowlers Gap), Garry Dowling (Fowlers Gap station manager), Mark Tilley, Derek Kennedy for their support during the project.

## Author contribution

ZB, SCG and STL conceived the study. ZB, KQ and STL collected data and ran analyses, ZB drafted the manuscript. All authors contributed to writing.

## Declarations

## Ethics approval

All sheep were treated using procedures approved by the University of New South Wales Animal Care and Ethics Committee in compliance with the Australian Code of Practice for the Use of Animals for Scientific Purposes (approval number 18/19B).

